# High School Science Fair: Ethnicity Trends in Student Participation and Experience

**DOI:** 10.1101/2021.12.03.471190

**Authors:** Frederick Grinnell, Simon Dalley, Joan Reisch

## Abstract

In this paper, we report ethnicity trends in student participation and experience in high school science and engineering fair (SEFs). SEF participation showed significant ethnic diversity. For survey students, the approximate distribution was Asian-32%; Black-11%; Hispanic-20%; White-33%; Other-3%. Comparing the SEF level at which students competed from school to district to region to state levels, we observed that black students made up only 4.5% of the students who participated in SEF beyond the school level, whereas students from other ethnic groups were more equally represented at all levels. The lower percentage of Black students resulted from a combination of lower overall participation in SEF and lower percentage of those students who did participate to advance to SEFs beyond the school level. Students who advanced to SEFs beyond the school level frequently received help from scientists, coaching for the interview, and were not required to participate in SEF. Black students received the least help from scientists, were least likely to receive coaching for the interview, and were most likely to be required to participate in SEF. They also were most likely to receive no help from parents, teachers, or scientists. Asian and Hispanic students (63.8% and 56.8%) indicated a greater interest in careers in science and engineering (S&E) compared to Black and White students (43.7% & 50.7%). In addition to career interest, the most important experiences that correlated with students who indicated that SEF increased their interests in S&E were getting help from the internet, books and magazines; getting help fine tuning the report; and overcoming obstacles by doing more background research, making a timeline, and perseverance. Black students did not report a positive effect of any of these strategies but experienced time pressure as more of an obstacle than did other students. Our findings identify a wide range of student experiences associated with positive SEF outcomes that could be enhanced for all students but especially Black students. More involvement of scientists in helping students who participate in SEFs would be particularly valuable.

## Introduction

Several years ago, we initiated a survey research program to learn student opinions about their experiences in high school science and engineering fairs (SEFs) with the goal of improving SEF practices. In this paper we report our latest findings based on an analysis of new questions regarding student ethnicity and level of SEF competition that we added to the surveys beginning in 2018.

SEFs represent something of an enigma. On one hand, they would seem to be an ideal means for students to experience for themselves the practices of science and engineering [1], now a central goal of Next Generation Science Standards (NGSS) [2]. Moreover, SEFs have become highly visible. In his 2011 State of the Union Address, President Obama remarked, “We need to teach our kids that it’s not just the winner of the Super Bowl who deserves to be celebrated, but the winner of the science fair” [3]. The film *Science Fair* was festival favorite at the 2018 Sundance Film Festival. Winners of the major SEFs can receive college scholarships worth $50,000 or more.

On the other hand, according to the National Center for Education High School Longitudinal Study of 2009/2012 (NCE-HSLS), only about 5% of the 23,500 high school students surveyed participated in a science competitions during high school [4]. Consistent with this finding, only about 5% of the almost 16,000 undergraduate students surveyed in the college Outreach Programs and Science Career Intentions survey (students in introductory freshman, mostly English, classes) reported participating in SEFs [5]. For many schools, the combination of budget limitations and competition for classroom time will decrease the possibility of promoting and supporting SEFs. Indeed, only 1/3 of the 900 schools surveyed in NCE-HSLS reported that they “hold math or science workshops or competitions” [4]. That number hasn’t changed much in the past 10 years judging from our recently reported survey of undergraduates (all on bioscience education trajectories) of whom about 60% attended high schools where participation in SEFs was not available [6].

SEFs began in the U.S. in New York City in the 1930’s. The stated goals were “to aid in the development of the scientific leaders of the next generation and at the same time foster a better understanding of science among its laymen” [7]. After the 1939–1940 World’s Fair in New York, SEF increasingly was viewed as a means to encourage and help students find their way to science and engineering career paths [8]. Using SEFs to accomplish STEM education for all students became an elusive goal.

When Microsoft Corporation carried out a STEM perceptions study, they found that only about 5% of the students said that they became interested in STEM because of SEFs [9]. Similarly, among students at a Queensland science and engineering university only 7% listed SEF as the reason they became interested in STEM [10]. Consistent with these reports, studies of national U.S. student cohorts suggest that the main effect of participating in high school science competitions is to help retain students who already are interested in STEM rather than attract those previously uninterested [5, 11]. Nonetheless, helping retain students already interested in STEM is an important outcome. Previously, we reported that 75-80% of high school students who participated in SEFs in 11^th^ and 12^th^ grades said that they were interested in a career in the sciences or engineering compared to just under 50% in 9^th^ grade. Moreover, we found that 6 out of every 10 undergraduates students we surveyed (all on bioscience education trajectories) had participated in SEF if they were available at the students’ high schools [6].

The question of how SEF outcomes vary in relationship to the type of SEF in which students engage has received little attention. Recent research on middle school SEFs identified three major types: mandatory SEF with high support (curriculum, class time, teacher engagement) (23% of students); mandatory with low support (57% of students); and voluntary with low support (20% of students) [12]. Previously, we reported that 60-70% of students who participated in high school SEFs were required to do so [13]. Students overwhelming disliked a competitive SEF requirement (4:1 or greater). They viewed competitive SEF as emphasizing winning rather than learning. Being required to compete reduced markedly the percentage of students who indicated that SEF increased their interests in the sciences and engineering (S&E) [14]. Since students uninterested in careers in S&E were most likely to participate in SEF in 9^th^ and 10^th^ grades, we suggested that schools should consider offering different SEF opportunities for students in 9^th^ and 10^th^ compared to 11^th^ and 12^th^ grades, e.g., a non-competitive rather than competitive option or incentivized rather than required [6, 15].

Recently, innovative high school programs have been described that combine student participation in SEF with student and teacher curricular support to promote STEM engagement and learning, especially focused on students from under-represented ethnic minorities and low socioeconomic backgrounds [16–18]. Such programs are particularly important given low levels of Black, Hispanic, and some other ethnic minority individuals in STEM fields compared with their proportion in the population. Underrepresentation occurs at the workforce level but begins in the education pipeline [19–21]. Trying to achieve diversification of the STEM workforce remains a major challenge and requires further development of creative programming at many levels including K-12 STEM education [22].

In this paper, we use our latest surveys to learn about ethnicity trends in student participation and experience in high school SEFs. SEF participation showed significant ethnic diversity. Overall, the data suggest that Black students are not getting the help that they need to be successful in SEFs compared to students in other ethnic groups. Our findings identify a wide range of student experiences associated with positive SEF outcomes that could be enhanced for all students, but especially for Black students. More involvement of scientists in helping students participating in SEFs would be especially valuable. Details are reported herein.

## Materials and methods

This study was approved by the UT Southwestern Medical Center IRB (#STU 072014-076). Study design entailed administering to students a voluntary and anonymous online survey using the REDCap survey and data management tool [23]. Survey recipients were high school students who participated in SEFs during the 2018/19 and 2019/20 school years using the Scienteer (www.scienteer.com) online portal adopted by Alabama, Louisiana, Maine, Missouri, Texas, Vermont, and Virginia for SEF registration, parental consent, and project management.

After giving consent for their students to participate in SEF, parents could consent for their students to take part in the SEF survey. However, to prevent any misunderstanding by parents or students about a possible impact of agreeing to participate or actually participating in the survey, access to the surveys was not available to students until after they finished <u>all</u> of their SEF competitions. When they initially signed up for SEF, students were told to log back in to Scienteer after completing the final SEF competition in which they participated. Those who did so were presented with a hyperlink to the SEF survey. Scienteer does not send out reminder emails, and no incentives were offered for remembering to sign back in and participate in the survey.

Survey content was the same as that used previously [13] except modified beginning in 2018 to include questions about student ethnicity and about the level of SEF competition in which students participated (school, district, regional, state). In 2019, a question was added about the location of the student’s high school (urban, suburban, rural). The survey can be found in supporting information (S1 Survey).

Survey data were summarized with frequency counts and percentages. Significance of potential relationships between data items was assessed using Chi-square contingency tables for independent groups. Results are presented two ways -- graphically to make overall trends easier to appreciate and in tables to show the actual numbers. A probability value of 0.05 or smaller was accepted as statistically significant but actual p values are shown. No adjustments were made for multiple comparisons.

## Results

### Overview of survey responses

Table 1 shows the number and ethnicity of almost 40,000 students/year who used the Scienteer website to participate in SEFs during 2018/19 and 2019/20. The table also shows the Scienteer students who completed SEF surveys. Scienteer students answer a question about their ethnicity at the time they register to participate in SEF. They self-identified most often as White (26%) and Hispanic (25%) followed by Asian (15%) and Black (7%). In addition, more than 25% of the students selected the Other category. Students who completed surveys were much less likely to choose the Other category (3%). For the survey students, the average percentages of Asian students (32%) and Black students (11%) were about twice as high as the original Scienteer students, slightly lower for Hispanic students (20%), and slightly higher for White students (33%).

**Table 1:** Number and ethnic distribution of students who signed up for SEF using Scienteer, Scienteer students who participated in the SEF survey, and overall U.S high school students.

| Data Set (# students) |  | Asian<br>% (#) | Black<br>% (#) | Hispanic<br>% (#) | White<br>% (#) | Other*<br>% (#) |
| --- | --- | --- | --- | --- | --- | --- |
| Scientist SEF students | 2018/19<br>(36,934) | 15.3<br>(5,662) | 6.8<br>(2,496) | 25.2<br>(9,304) | 26.2<br>(9,690) | 26.5<br>(9,782) |
|  | 2019/20<br>(38,570) | 14.6<br>(5,365) | 6.6<br>(2,504) | 24.5<br>(9,207) | 26.1<br>(10,010) | 28.1<br>(11,484) |
| Scientist students who<br>completed surveys | 2018/19<br>(382) | 28.8<br>(110) | 13.1<br>(50) | 22.8<br>(87) | 32.2<br>(123) | 1.8<br>(7) |
|  | 2019/20<br>(938) | 35.0<br>(328) | 9.1<br>(85) | 17.4<br>(163) | 33.5<br>(314) | 4.1<br>(38) |
| 2009 High School NCES<br>X2RACE Ethnicity Profile | 2012<br>(23,503) | 3.6<br>(828) | 13.6<br>(3,128) | 22.1<br>(5,083) | 51.9<br>(11,937) | 8.9<br>(2,047) |

\*Includes American Indian; Hawaiian Native/Pacific Islander; and various mixed-race combinations.

In relationship to U.S. high school data shown in the last row of Table 1 [24], obvious differences in ethnicity between SEF/survey students and the overall high school population were overrepresentation of Asian students and underrepresentation of White students. We treat the Scienteer SEF population as a national group of high school students. However, it should be recognized that these students may not be truly representative of a national sample since they come from only 7 U.S. states, and they only attend high schools where SEFs are available.

Overall, 1-2% of all students who signed up for SEFs through Scienteer completed surveys, consistent with our previously reported experience with 2016/17 and 2017/18 Scienteer cohorts [14]. Given that student participation in the surveys involves an indirect, single electronic invitation without incentive or follow-up, the low response rate has not been surprising [25–27]. Most of the submitted surveys (greater than 92%) were complete and non-duplicates. The completed surveys were used for data analyses, and the data sets can be found in supporting information S1 Dataset and S2 Dataset for 2018/19 and 2019/20 school years respectively.

Previously, we found that student survey answers were reproducible from year to year [14]. That trend continued for the 382 students in 2018/19 and 938 students in 2019/20. Supplemental Table 1 (S1 Table) shows results for the entire set of 100+ possible survey questions and answers regarding student demographics, opinions about SEF, help received, obstacles encountered, and ways of overcoming obstacles. Answers selected by the 2018/19 and 2019/20 cohorts not only were similar to each other, but also were similar to the combined 2016/17 and 2017/18 results.

Fig 1 shows results for selected features from the 2018/19 and 2019/20 surveys that, along with student ethnicity, will be the major focus of this paper. About 2/3 of the students who completed surveys said that they were required to participate in SEF. About half received help doing SEF from parents and/or teachers but less than 10% from scientists. An important point that we realized in analyzing the new SEF surveys, which we had not noticed previously, is that a quarter of the students did not receive help from parents, teachers, or scientists.

**Fig 1:**
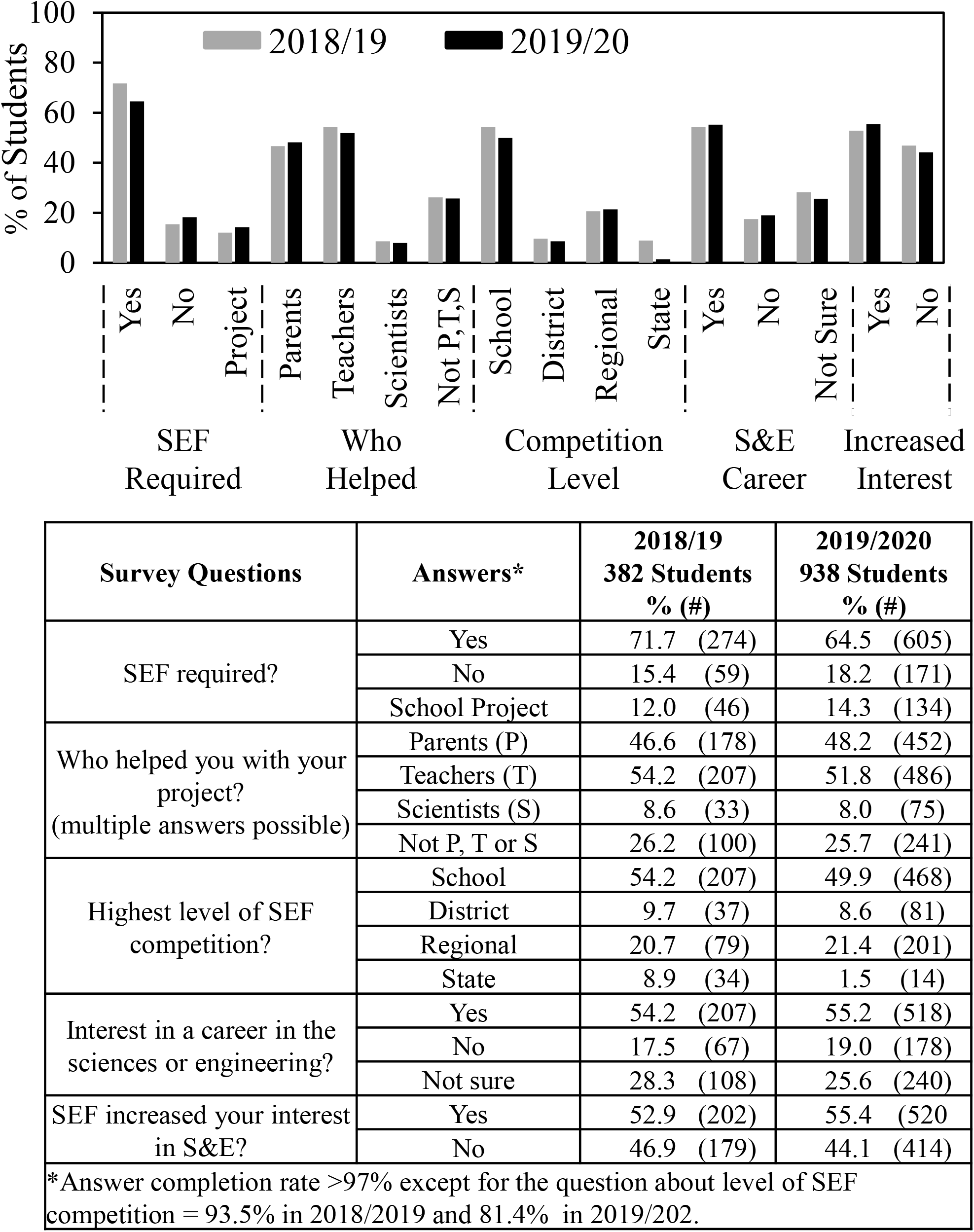
Some key features of SEF experience -- 2018/19 vs. 2019/20 student groups.

In addition to the question about ethnicity, beginning in 2018/19 we also began asking about the level of SEF in which students participated. About half the students participated in SEF at the school level; the remainder went on to higher level SEFs. The lower percentage of students who competed in state level SEF in 2019/20 resulted from changes in SEF opportunities in late spring 2020 because of COVID-19 restrictions. The completion rate for almost all the survey questions was >97% except for the question about level of SEF competition.

Finally, Fig 1 shows that of the students who completed surveys in 2018/19 and 2019/20, slightly more than half said that they were interested in a career in the sciences and engineering; 20% were uninterested; and the remainder unsure. Also, slightly more than half the students said that participating in SEF increased their interest in S&E.

### Students’ ethnicity and SEF experience

For subsequent figures, the 2018/19 and 2019/20 student survey results were combined. Fig 2 shows ethnicity demographic data. Most students did SEFs in 9^th^ and 10^th^grades. However, the proportion of the Hispanic students doing SEFs in 11^th^ and 12^th^ grades (33%) was higher than for other groups – white (21%), Asian (17%), black (23%). More females than males completed surveys; the difference was more than 2:1 for Black students. Most (80%) of White, Asian and Black students who completed surveys attended suburban schools, whereas the Hispanic student population was more evenly distributed with 50% in suburban schools and just under 40% in urban schools.

**Fig 2.**
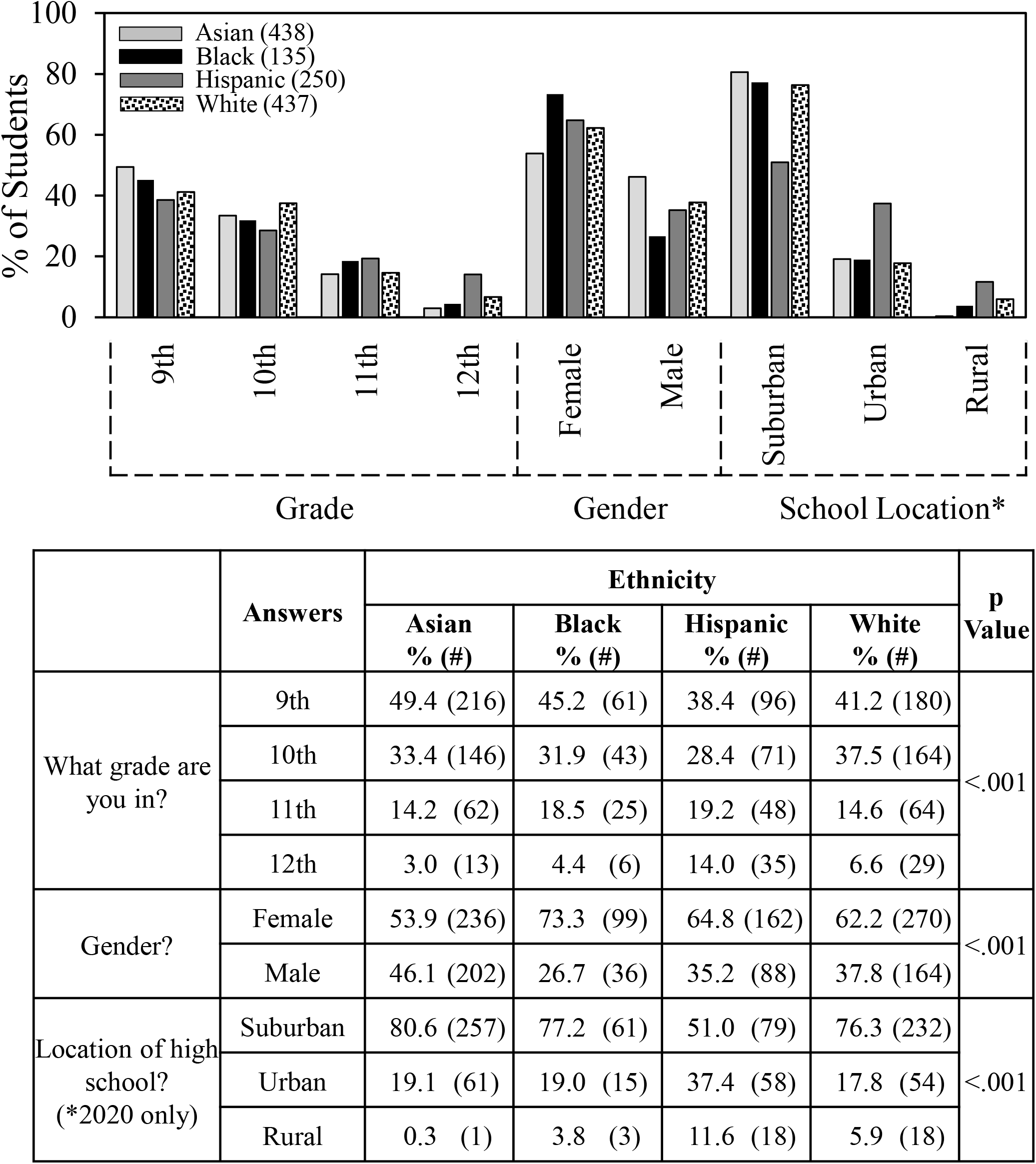
Ethnicity and student demographics.

Fig 3 shows data about student involvement. The requirement to participate in SEF ranged from 84% for Black students to 58% for Asian students. About half the students in each ethnic group reported help from parents with White students receiving the most help (54%). Also, about half the students in each group received help from teachers. In response to the paired questions, ‘did you get the amount of help you wanted from teachers’ and ‘did you get the kind of help that you wanted from teachers,’ around 70% of the students responded “yes” to each question.

**Fig 3.**
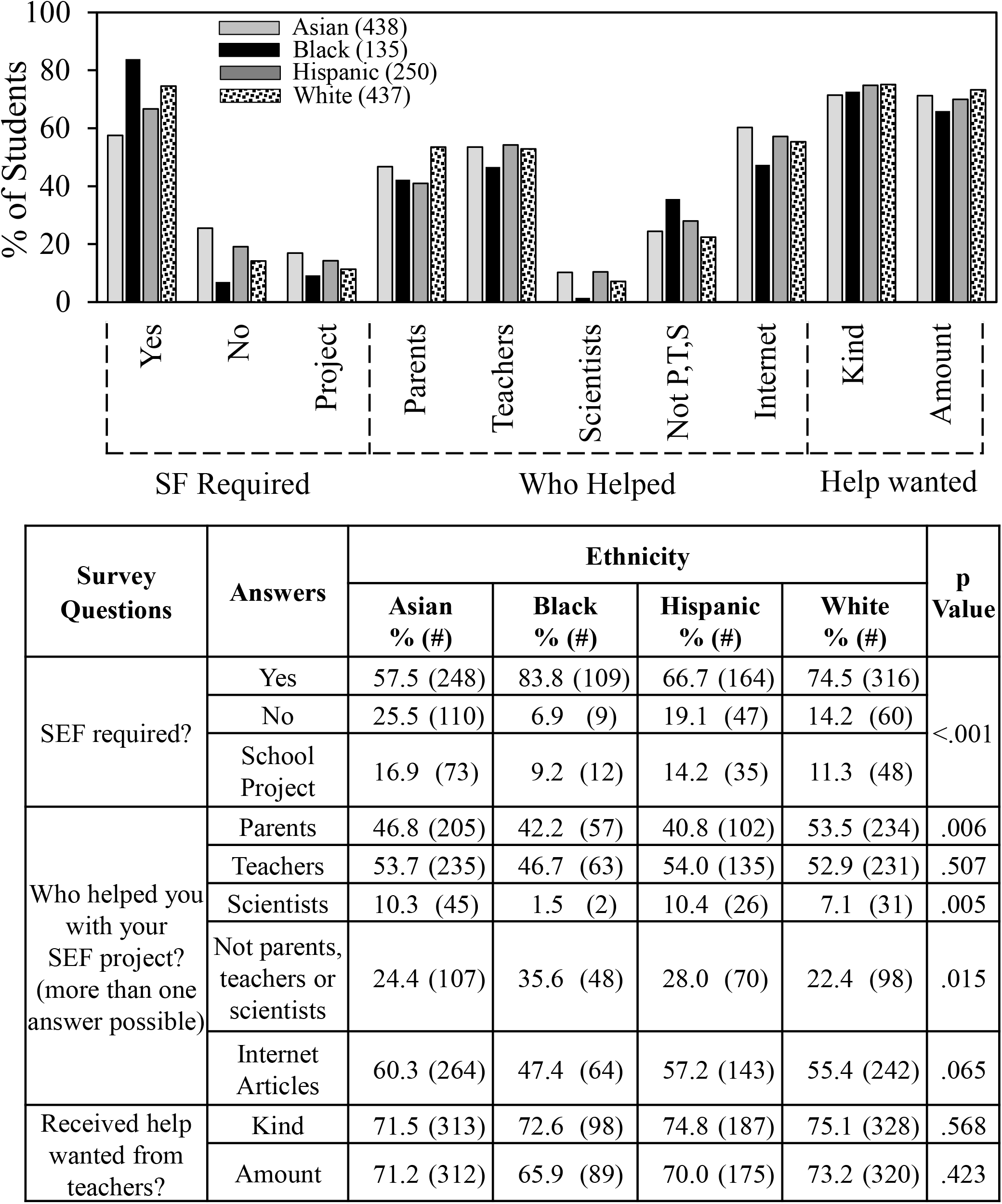
Ethnicity and SEF involvement.

Black students received much less help from scientists compared to students in the other ethnic groups, i.e., 2% of the Black students vs. 10% of the Asian and Hispanic students and 7% of the White students. Also, a higher percentage of Black students (36%) compared to students in the other groups (22-28%) reported receiving no help from parents, teachers, or scientists. For all ethnic groups, students received more help from internet articles than from other sources.

Scienteer tracks whether students participate in SEF at only the school level or move on to more advanced SEFs. Table 2 shows that almost 25% of the Scienteer students advanced to SEFs beyond the school level. That number was higher (40%) for students who completed surveys. Both Scienteer and survey data show that Black students made up less than 5% of all students who reached more advanced SEFs. Hispanic and White students were present in advanced SEFs in roughly equal numbers (28%). Asian and Other students were present at 17% and 23% respectively according to Scienteer data but 37% and 1% according to survey data.

**Table 2:**
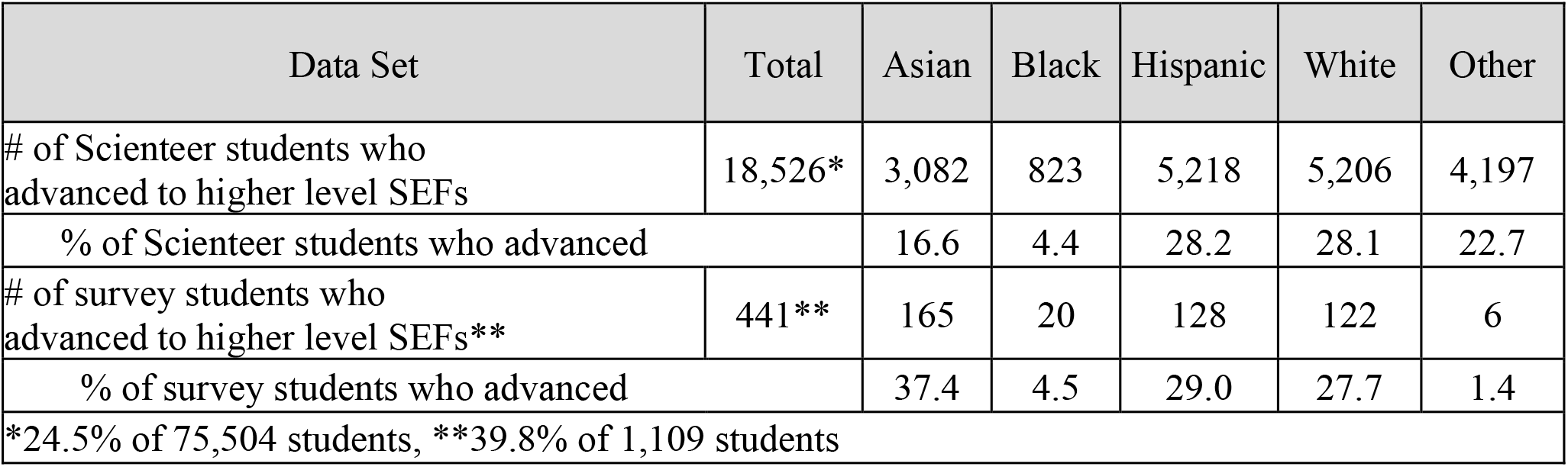
Distribution by ethnicity of 2018/19 and 2019/20 Scienteer and survey students who competed in higher level SEFs (district, region, or state)

| Data Set | Total | Asian | Black | Hispanic | White | Other |
| --- | --- | --- | --- | --- | --- | --- |
| # of Scienteer students who advanced to higher level SEFs | 18,526* | 3,082 | 823 | 5,218 | 5,206 | 4,197 |
| % of Scienteer students who advanced |  | 16.6 | 4.4 | 28.2 | 28.1 | 22.7 |
| # of survey students who advanced to higher level SEFs** | 441** | 165 | 20 | 128 | 122 | 6 |
| % of survey students who advanced |  | 37.4 | 4.5 | 29.0 | 27.7 | 1.4 |
| *24.5% of 75,504 students, **39.8% of 1,109 students |  |  |  |  |  |  |

Fig 4 summarizes in more detail data about SEF outcomes based on the surveys. About 80% of the Black students surveyed only competed in SEF at the school level compared to 67% of the White students, 57% of the Asian students, and 41% of the Hispanic students. Comparing different groups, Hispanic students were most likely percentagewise to compete in district, regional and state level SEFs. Fig 4 also shows that Asian and Hispanic students reported a greater interest in careers in science and engineering (64% and 57% respectively) compared to White and Black students (51% and 44%) and were more likely to indicate that participating in SEF increased their interest, 64% for Asian and Hispanic students vs. 46% for White and Black students.

**Fig 4.**
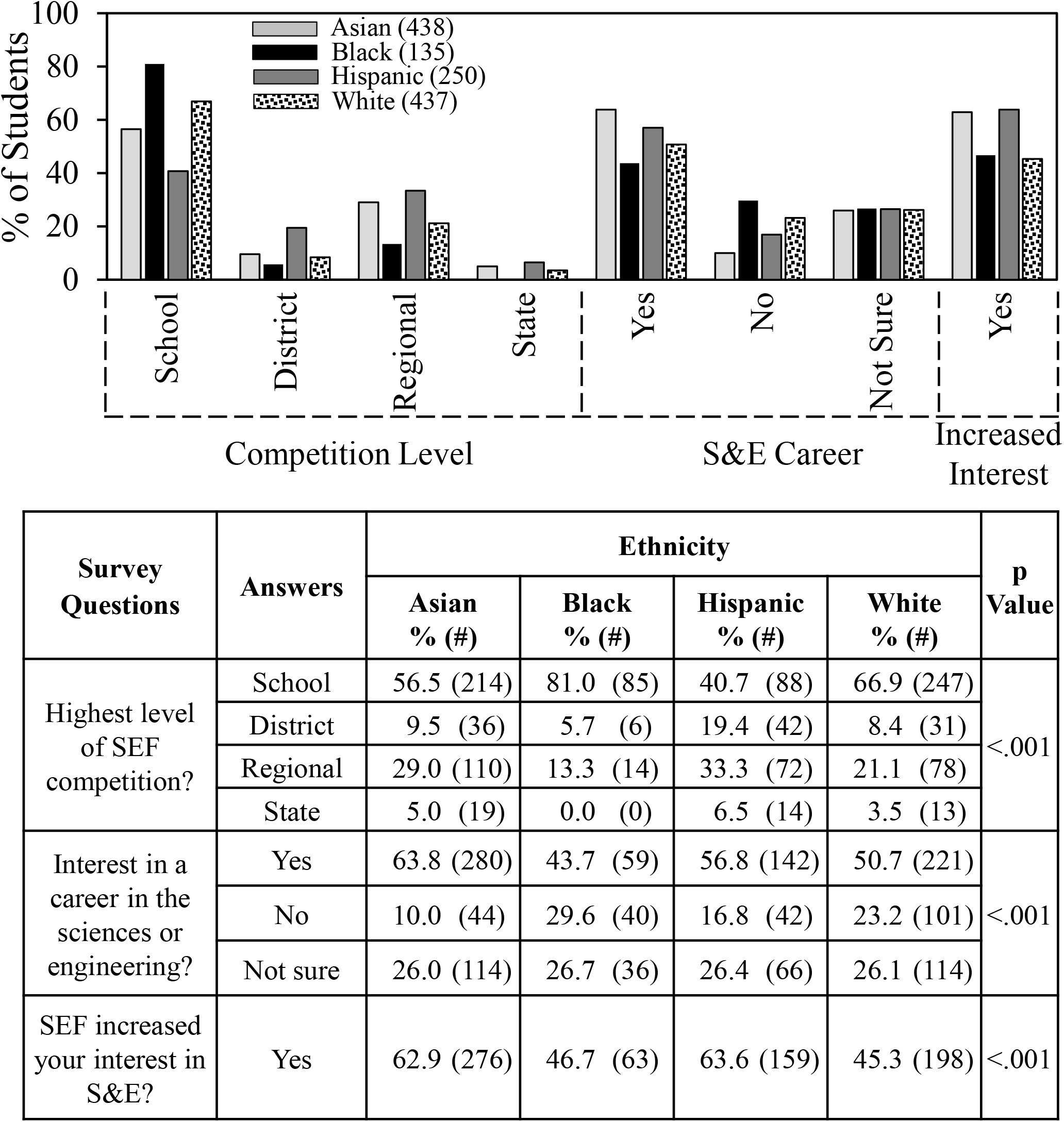
Ethnicity and SEF outcomes.

### Differences in students’ experiences associated with sources of help and level of SEF competition

Another way to analyze the results was by comparing student experience according to from whom the students received help. Fig 5 compares the consequences of receiving help from scientists (which could include teachers and/or parents); parents or teachers but not scientists; and no help from parents, teachers, or scientists. 108 students received help from scientists. More than 80% of those students participated in SEFs beyond the school level. By comparison, 871 students had help from parents and or teachers but not scientists. About half those students did SEF at the school level and the other half advanced to higher level SEFs. Students having help from scientists also was associated with an increase in the likelihood that students would indicate they were interested in a career in the sciences or engineering and an increase in the likelihood that students would indicate that SEF increased their interest in the sciences or engineering. Help from parents and/or teachers had less impact than help from scientists but more so than no help from scientists, teachers or parents.

**Fig 5.**
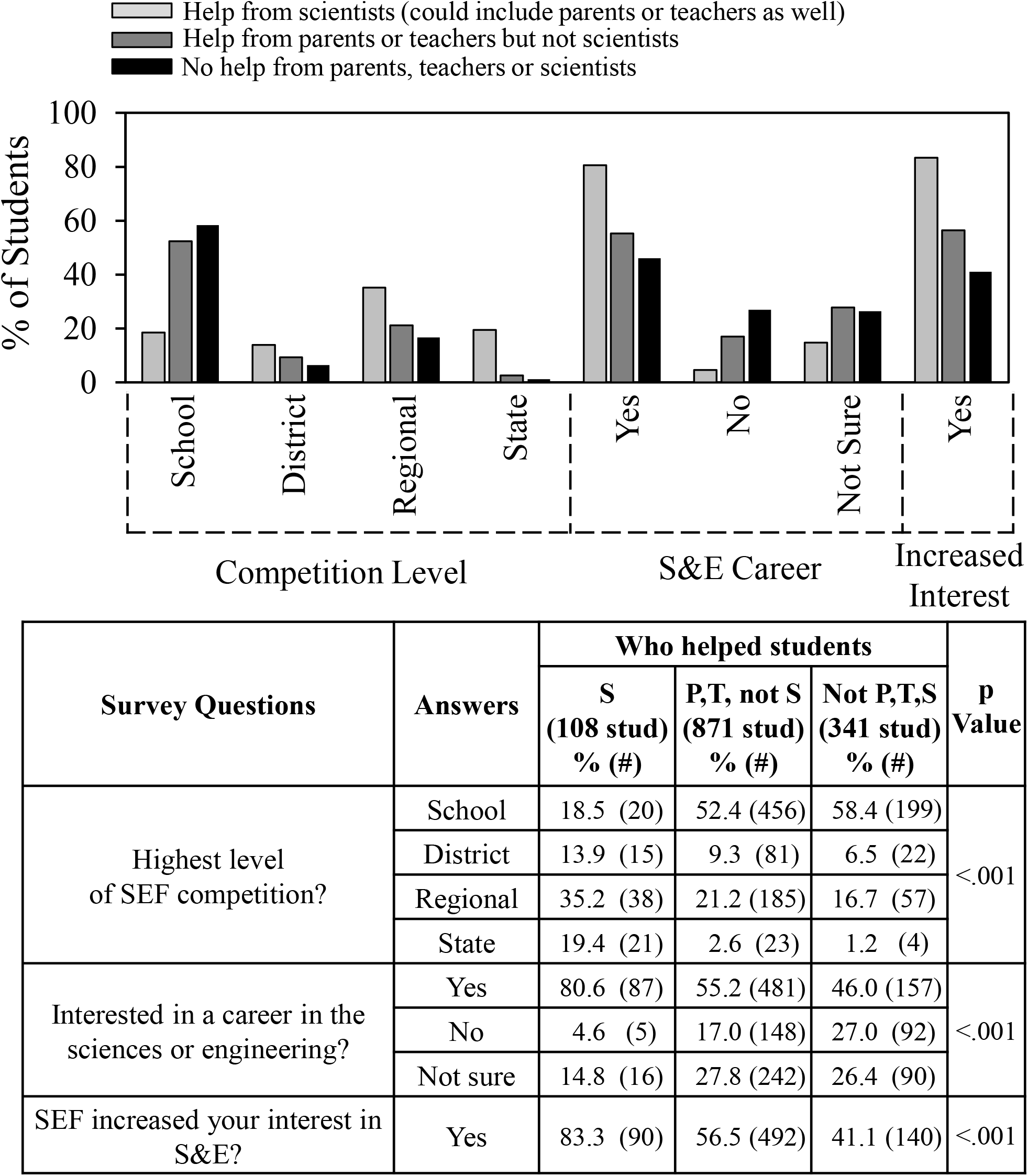
Impact of help from scientists, parents and teachers.

We also compared differences in student experience according to whether students competed only at the school level or in more advanced SEFs. Fig 6 shows even more clearly than Fig 4 that Black students were the least likely to advance beyond school level SEFs. By comparison, of the Hispanic students who indicated the level of SEF in which they participated, 2/3 went beyond the school level. By the criterion of whether they advanced to SEFs beyond the school level, Hispanic students were more successful in SEF than students in other ethnic groups. Students who advanced to higher level SEFs also were more likely to have an interest in a career in the sciences or engineering; almost twice as likely to indicate that SEF participation increased their interest in S&E; and less likely to have been required to do SEF.

**Fig 6.**
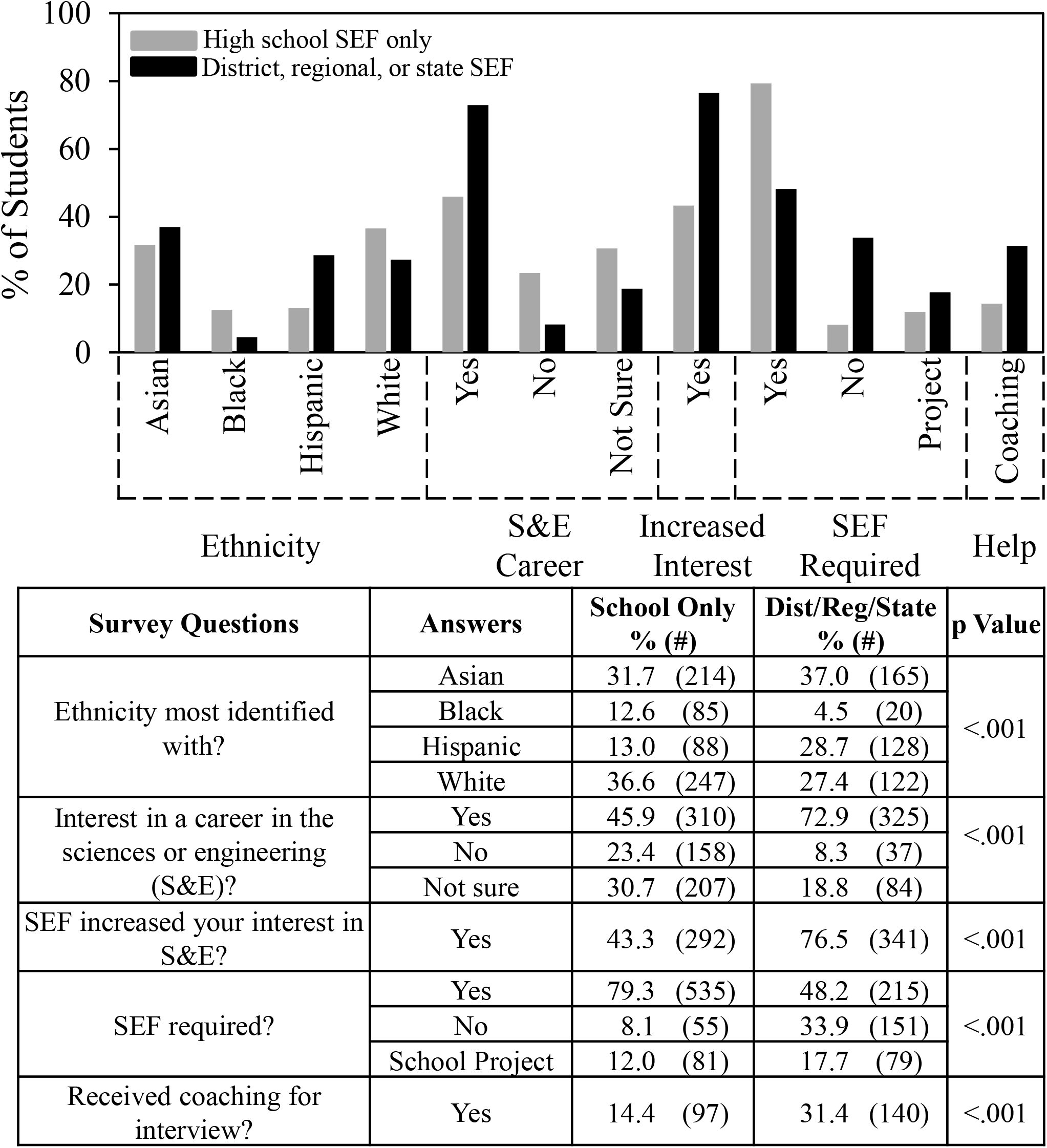
Level of SEF competition.

One type of help that students received stood out compared to all others. Students who said that they received coaching for the SEF interview were more than twice as likely to advance beyond the school level compared to students who did not receive coaching.

### Factors affecting impact of SEF on student interest in the sciences and engineering

One of the most important goals of SEFs is to increase student interest in S&E. As was shown in Fig 4, Hispanic and Asian students were more likely to indicate that SEF increased their interest in S&E compared to Black and White students. Figs 7, 8 and 10 show the survey data analyzed to determine what experiences by students in each ethnic group correlated with a greater percentage of students who indicated that SEF increased their interest in S&E. Some of these features turned out to be typical of students in all ethnic groups, while others were more selective. Fig 7 concerns overall features. Fig 8 concerns sources of help. Fig 10 concerns types of help received, obstacles faced, and ways to overcome obstacles. Numerical and statistical details for the figures can be found organized by ethnic group in supplemental data (S2–S5 Tables).

**Fig 7.**
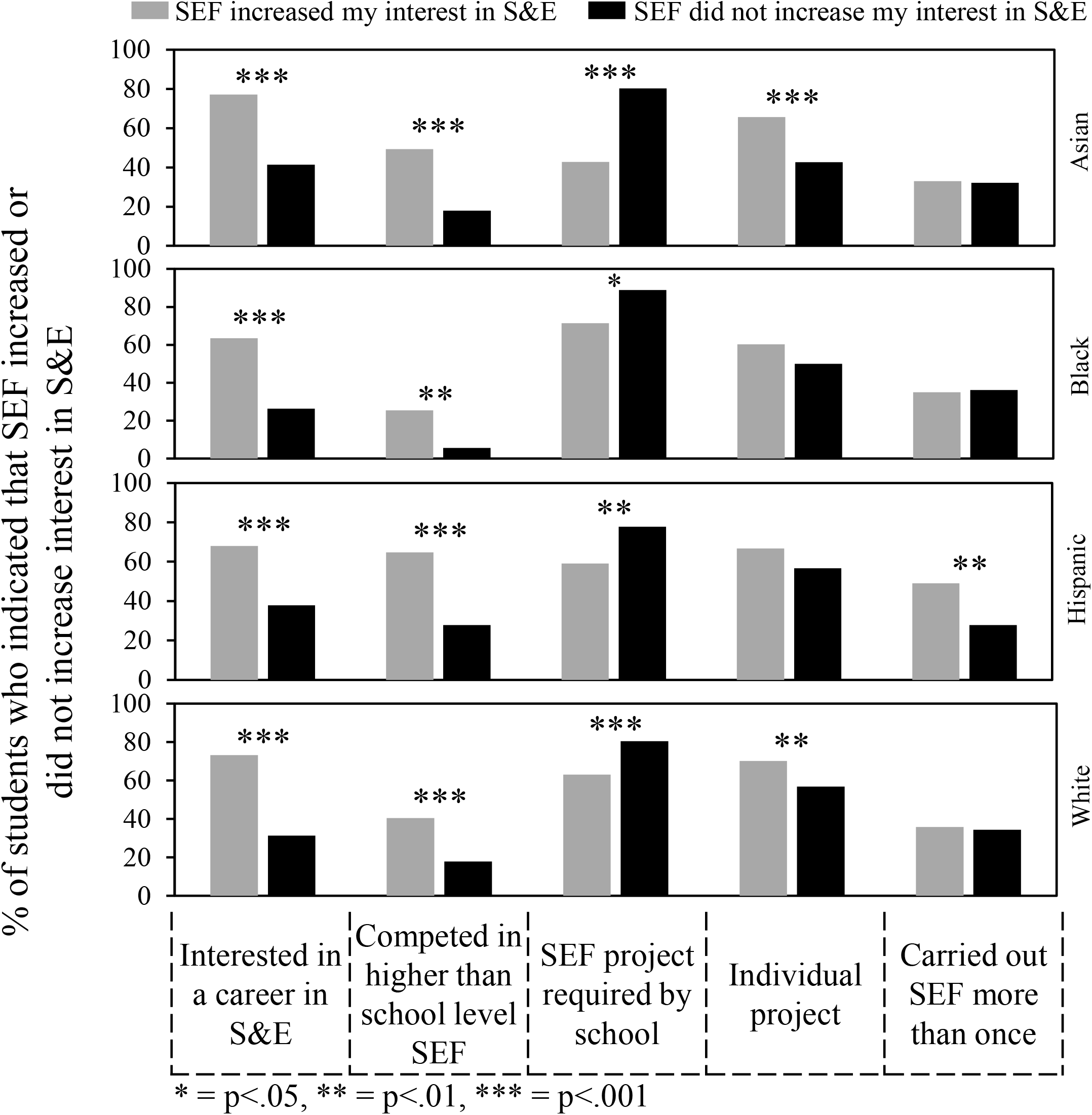
Overall features that influenced impact of SEF on student interest in S&E.

**Fig 8.**
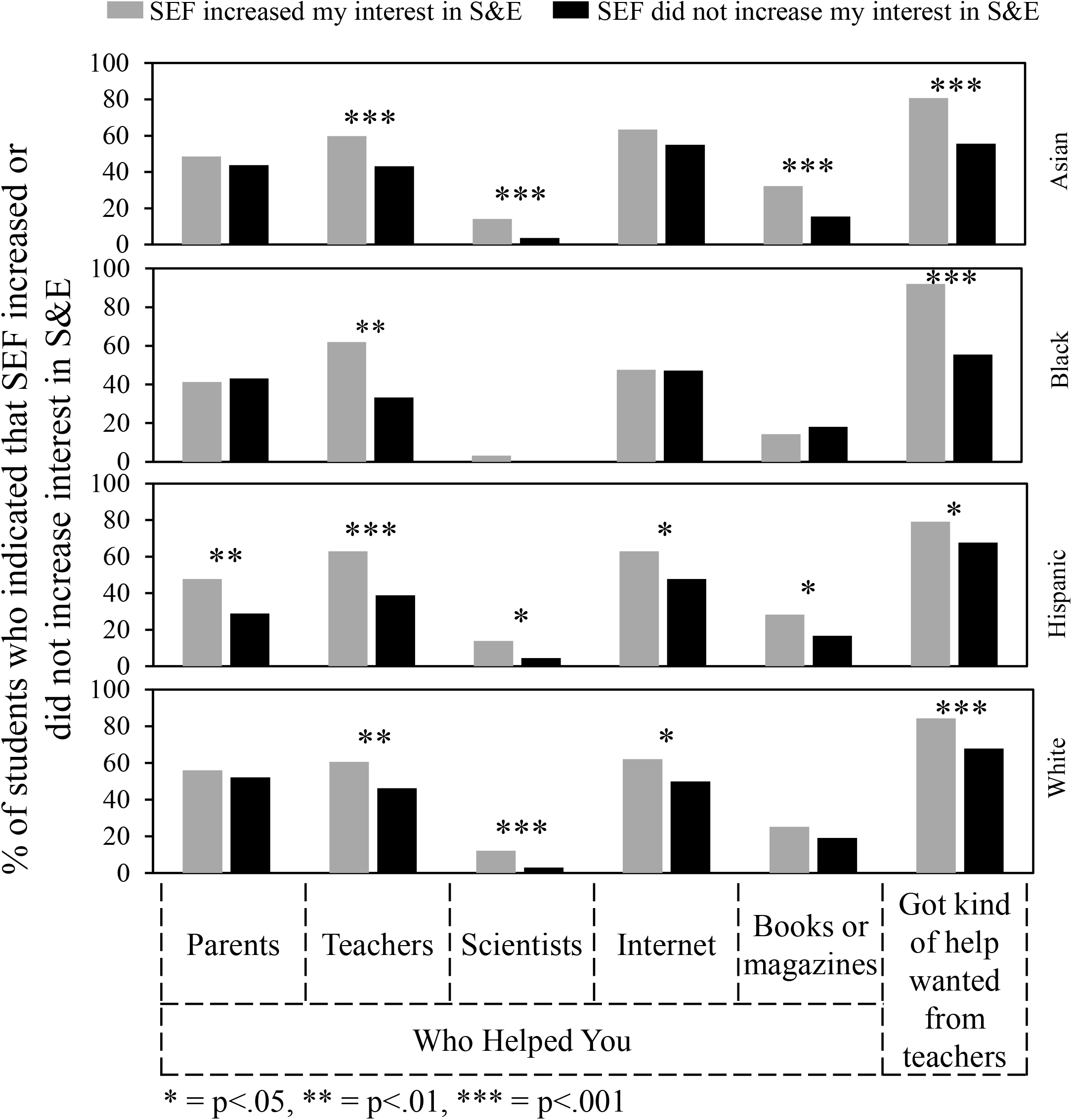
Sources of help that influenced impact of SEF on student interest in S&E.

**Fig 9.**
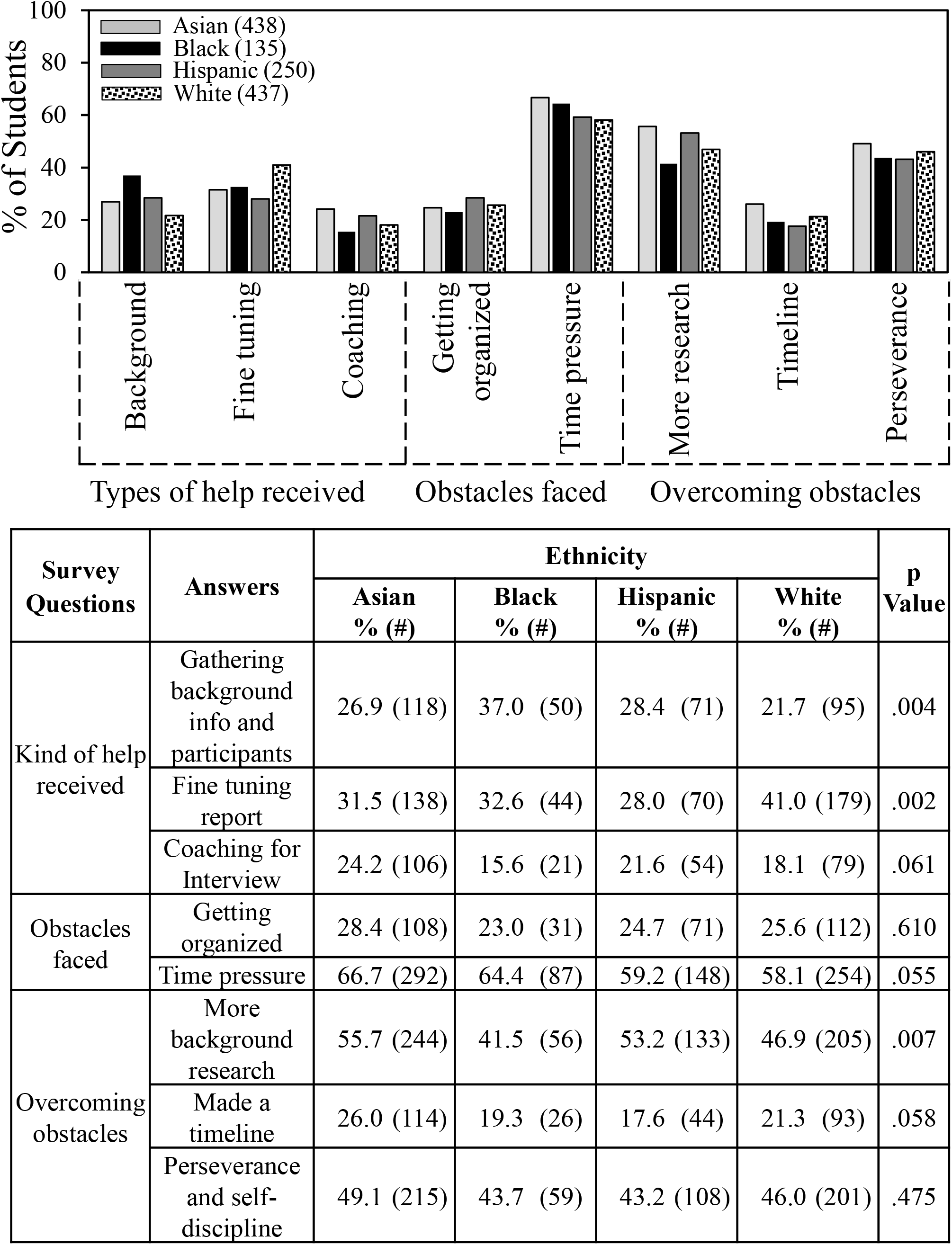
Ethnicity and SEF help and obstacles.

**Figure 10.**
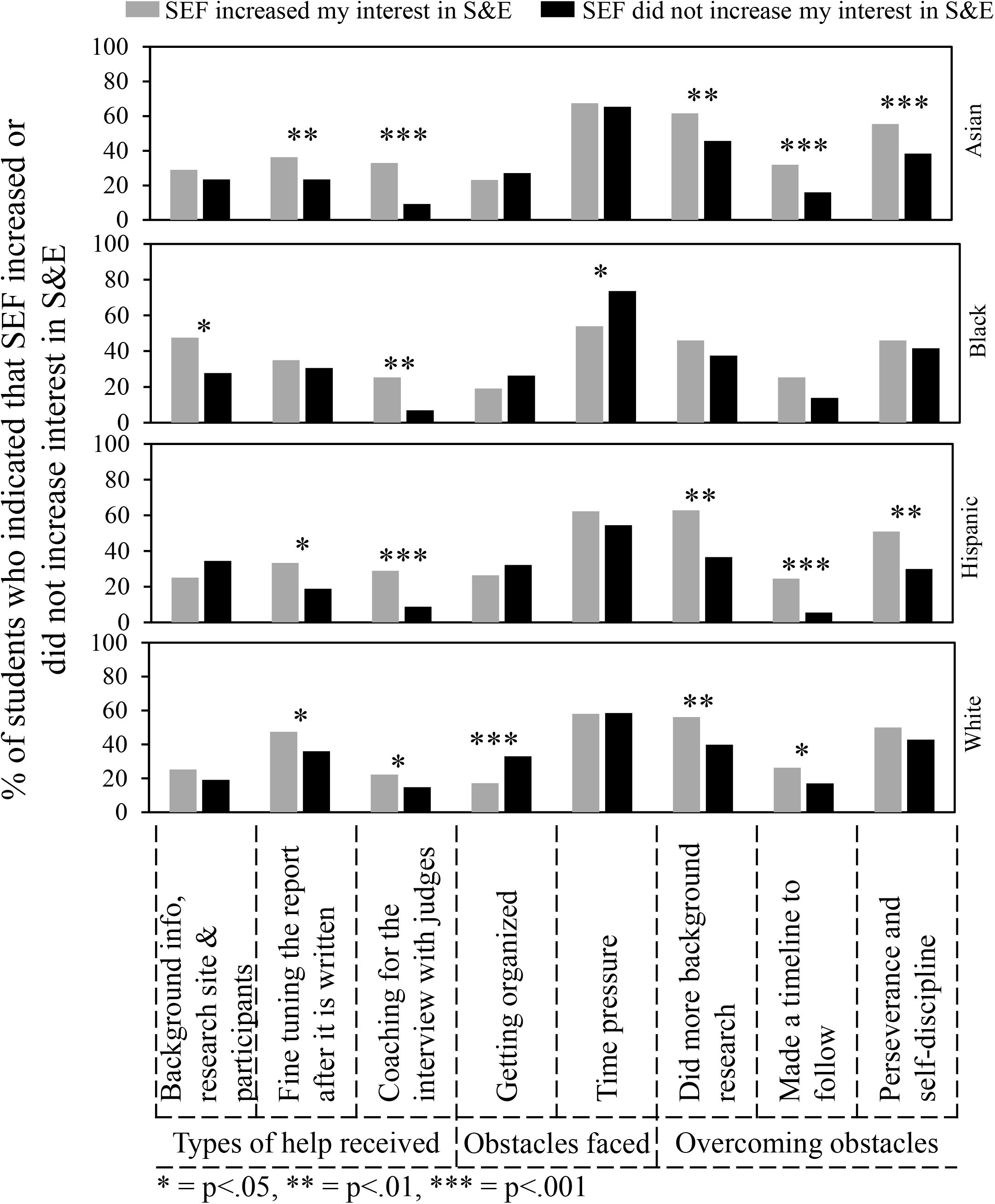
Kinds of help, obstacles faced, and ways to overcome obstacles that influenced impact of SEF on student interest in S&E.

Fig 7 shows that students in all ethnic groups who indicated that SEF increased their interests in S&E were more likely be interested in a career in S&E compared to students who indicated that SEF did not increase their interest in S&E. Numerical values for this comparison can be found in row #1 of supplemental Tables S2–5: Asian students, 77.2% vs. 41.4% (p<.001); Black students, 63.5% vs. 26.4% (p<.001); Hispanic students, 67.9% vs. 37.8% (p<.001); White students, 73.2% vs. 31.4% (p<.001). Also, students in all ethnic groups who said that SEF increased their interests in S&E were more likely to indicate that they participated in SEF beyond the school level. On the other hand, if they were required to participate in SEF, then they were more likely to indicate the opposite. Carrying out an individual rather than team project was more positive for Asian and White students. Carrying out SEF more than once was more positive for Hispanic students.

Fig 8 is organized similarly to Fig 7. For instance, getting help from parents had a positive impact for Hispanic students who said that SEF increased their interests in S&E. Help for parents did not make a difference for students in the other ethnic groups. Numerical values for this comparison can be found row #6 of supplemental Tables S2–5 -- Asian students, 48.6% vs. 43.8% (p=.338); Black students, 41.3% vs. 43.1% (p=.860); Hispanic students, 47.8% vs. 28.9% (p=.005); and White students, 56.7% vs. 52.1% (p=.411). Also, for students in all groups, getting help from teachers and getting the kind of help wanted had a positive impact on students who indicated that SEF increased their interests in S&E. Help from scientists was positive for students in all the groups but effectively unavailable to Black students as shown earlier (Fig 3 and Fig 5). Hispanic, Asian, and White students but not Black students also indicated a positive impact of some combination of using articles from the internet and books or magazines.

More subtle differences than those shown in Figs 7 and 8 occurred regarding kinds of help students received, obstacles faced, and ways to overcome obstacles. Fig 9 summarizes the most important factors, which showed only small differences between ethnic groups. Black students appeared to receive more help that others gathering background information/logistics and were less likely to do more research to overcome obstacles. White students appeared to receive more help fine tuning their reports. Although experienced similarly overall by students in the different ethnic groups, the factors shown in Fig 9 turned out to be differentially important with respect to the impact of SEF on student interest in S&E.

Fig 10 shows that getting help with background information/logistics made more of a difference for Black students compared to other students in relationship to whether students indicated that SEF participation increased their interests in S&E. Numerical values for this comparison can be found in row #12 of supplemental Tables S2–5: Asian students, 29.0% vs. 23.5% (p=.208); Black students, 47.6% vs. 27.8% (p=.021); Hispanic students, 25.5% vs. 34.4% (p=.144); White students, 25.3 vs. 19.1% (p=.121). For students in all ethnic groups, coaching for the interview had a positive impact on the percentage of students who indicated that SEF increased their interests in S&E. Although time pressure was the most frequently indicated obstacle faced, the negative impact appeared to be greater for Black students than others. Getting help fine tuning their reports benefitted students in other groups more than Black students. Getting organized was an obstacle for White students. Hispanic and Asian students (and to a lesser extent White students) indicated a positive impact of overcoming obstacles by doing more background research, making a timeline to follow, and perseverance. None of these ways of overcoming obstacles was statistically significant for Black students.

## Discussion

In this paper, we report ethnicity trends in student participation and experience in high school science and engineering fairs. Our findings are based on students who registered and carried out SEF projects through the Scienteer website (almost 80,000 students over two years) and on those students who completed surveys after finishing all of their SEF competitions (1,320 students over two years). We treat the Scienteer SEF population as a national group of high school students. However, it should be recognized that these students may not be truly representative of a national sample since they come from only 7 U.S. states, and they only attend high schools where SEF is available.

SEF participation showed significant ethnic diversity. According to Scienteer registration, the approximate distribution of students was Asian-15%, Black-7%, Hispanic-25%, White-26%, and Other-27%. For survey students, the approximate distribution was Asian-32%, Black-11%, Hispanic-20%, White-33%, and Other-3%. Comparing the survey results to the original Scienteer registration, twice as many students self-identified as Asian, almost twice as many as Black, and very few selected Other. We don’t know why the difference in selection of the Other category. One possibility is that during initial Scienteer registration some students preferred to have ethnic anonymity, which was less important to them after completing all of their SEF activities.

The national NCE-HSLS (2009) study determined the percentage of students who indicated that they planned to pursue a STEM major when they reached college [28]. Broken down by ethnic identification, the STEM-interested student groups were Asian (41.9%), Black (15.5%), Hispanic (19.8%), and White (24.8%). Under representation of Black and Hispanic individuals in STEM careers does not result from an initial lack of student interest in STEM but from subsequent differences in persistence and overall graduation rates in STEM degree programs compared to Asian and White students [29, 30].

The pattern of students who participated in SEF resembles the distribution of STEM-interested student groups. An attractive hypothesis is that even though only a small portion of high school students participate, those who do are representative of overall STEM interests amongst high school students. Participating in high school science competitions helps retain students’ interest in STEM [5, 11]; SEFs provide an opportunity to practice those interests [6]; and authentic research experiences help high school students understand the nature of science [31]. Therefore, increasing the availability of SEFs to more students would be a valuable objective.

Based on Scienteer and survey data, Black students made up only 4.5% of the students who participated in SEFs beyond the school level, whereas students from other ethnic groups were more equally represented. The lower level could result from two different factors -- on one hand the overall lower level of participation of Black students in SEFs compared to other ethnic groups; and on the other hand, the lower success rate of those black students who did participate to advance beyond the school level. As will be discussed, the data suggest that Black students are not getting the help that they need to be successful in SEFs.

The ratios (percentages) for experiences that appeared to be especially relevant to whether students advanced to SEFs beyond school only were 3.7:1 for getting help from scientists; 2.2:1 for being coached for the interview; 1.7:1 for not being required to participate in SEF; and 1.6:1 for being interested in a career in S&E. On the other hand, the ratio was 0.42:1 for getting no help from parents, teachers, or scientists. Several of these factors likely contribute to the lower percentage of Black students that advanced. Black students received the least help from scientists; were least likely to get coaching for the interview; most likely to be required to do SEF; were least interested in a career in S&E; and were most likely to get no help from parents, teachers or scientists. Hispanic students were the most likely to advance beyond school level SEFs followed by Asian and White students. A limitation of our analysis is the uncontrolled factor that the number of advancing students will depend not only on student performance, but also on the number of SEF entries that a school or district permits to advance, which typically will be governed by individual school and district policies.

The question regarding level of SEF competition was new to the surveys analyzed in this report. However, the findings are consistent with our previous analysis of a cohort of high school students from a North Texas school district whose students are consistently high achievers in the Dallas Regional Science and Engineering Fair and beyond. We found that 26% of those students received help from scientists; over 80% received coaching for the interview; and less than 10% were required to participate in SEF [32].

One of the most important goals of SEF is to increase student interest in S&E. Asian and Hispanic students indicated that SEF participation increased their interests in S&E (63.6% and 63%), which was higher than Black and White students (46.7% and 45.3%). These differences were similar to those indicated by students about their interests in STEM careers -- Asian and Hispanic students (63.8% and 56.8%) vs. Black and White students (43.7% & 50.7%). Therefore, students already interested in S&E careers might be primed to have a positive SEF experience. Or participation in SEF may consolidate students’ interest in STEM careers, consistent with the findings about retention via science competitions cited earlier [5, 11].

Other student experiences that potentially enhanced SEF increasing a student’s interest in S&E were competition beyond school level; help from teachers; and coaching for the interview. A requirement to participate was negative. For Asian, Hispanic, and White students, help from scientists (not available to Black students) was positive. Also, help from the internet and/or books and magazines was positive for Asian, Hispanic and White students but made no difference for Black students.

Only Hispanic students indicated that help from parents was positive. Also, only Hispanic students indicated that carrying out SEF more than once was positive. The latter was consistent with the observation that a higher percentage of Hispanic students participated in SEF in 11^th^ and 12^th^ grades. Previously, we reported that high school students who participated in SEF in 11^th^ and 12^th^ grades were more likely to indicate that they were interested in a career in S&E, and more than 50% of undergraduates we surveyed (all on bioscience education trajectories) who participated in SEFs in high school did so more than once [6].

Getting help with background information/logistics was more positive for Black students than others. Time pressure was more of an obstacle for them. Also, Black students did not experience a positive effect of fine tuning the report. Getting organized was more of an obstacle for White students than others. The ways of overcoming obstacles that many students indicated had a positive effect were doing more background research, making a timeline, and perseverance. Black students did not report a positive effect of any of these ways to overcome obstacles.

These findings identify a wide range of student experiences associated with positive SEF outcomes that could be enhanced for all students, but especially for Black students. More involvement of scientists in helping SEF students would be particularly valuable since 80% of students who received help from scientists said that they were interested in a career in S&E; less than 5% said that they were uninterested; and 83% of the students said that help from scientists increased the positive impact of participating in SEF.

Previous studies by others showed factors contributing to success in SEF included parental support and encouragement [33], social and research resources [34], and access to outside of school facilities [35]. In our work, we observed that students who received help from scientists had an easier time getting their research idea, more access to articles in books and magazines, and less difficulty getting the resources to carry out the projects [13]. SEF guidelines typically emphasize that a SEF project should be the student’s; the mentor’s job is supportive not directive. Nevertheless, because help from scientists and similar support is available to only a subset of students, some critics have suggested that competitive SEF is fundamentally unfair [36–38]. For this reason, we suggested the possible value of offering students a non-competitive SEF option [15]. For competitive SEF, our findings support the idea that the best way to increase success of all high school students would be by developing programs that result in more help for the students from scientists or student scientists. Some innovative programs along these lines recently have been initiated [16–18, 39].

## Acknowledgments

We are grateful to Russell Cowen and Rocky Slavin, managers of Scienteer Technologies, who incorporated the parental consent and SEF survey REDCap links into the Scienteer website and continue to provide ongoing oversight and management of survey access. Dr. Elise Christopher from the National Center for Education Statistics advised us about the *High School Longitudinal Study of 2009*.

## Supporting information

S1 Data set. Excel dataset showing all of the survey questions and answers for 2018/19 surveys.

S2 Data set. Excel dataset showing all of the survey questions and answers for 2019/20 surveys.

S1 Survey. Survey questions.

**Supplemental Table 1.** Comparison of survey responses by students from 2018-19 vs. 2019-2020 vs. combined 2016-18 survey years

| Survey Questions | Answers | 2018/19<br>(382<br>students) | 2019/20<br>(938<br>students) | 2016-18<br>(363<br>students) |
| --- | --- | --- | --- | --- |
|  |  | % Students |  |  |
| 1. What grade are you in? | 9th | 44.8 | 43.6 | 40.5 |
|  | 10th | 31.9 | 33.8 | 32.2 |
|  | 11th | 13.9 | 16.5 | 11.3 |
|  | 12th | 8.6 | 5.8 | 11.0 |
| 1A. Location of high school? | Suburban | - | 70.5 | - |
|  | Urban | - | 20.7 | - |
|  | Rural | - | 4.8 | - |
| 2. Gender? | Female | 63.9 | 59.3 | 63.4 |
|  | Male | 34.3 | 40.2 | 35.8 |
| 2A. Ethnicity most identified with? | Asian | 28.8 | 35.0 | - |
|  | Black | 13.1 | 9.1 | - |
|  | Hispanic | 22.8 | 17.4 | - |
|  | White | 32.2 | 33.5 | - |
|  | Other | 1.8 | 4.1 | - |
| 3. During high school have you carried out science fair more than once? | Yes | 29.6 | 37.5 | 31.1 |
|  | No | 69.4 | 61.8 | 67.2 |
| 3A. In which science fair competitions did you compete this year? | School | 54.2 | 49.9 | - |
|  | District | 9.7 | 8.6 | - |
|  | Regional | 20.7 | 21.4 | - |
|  | State | 8.4 | 1.5 | - |
| 4. Was your science fair project Team or Individual? | Individual | 63.1 | 58.6 | 70.2 |
|  | Team | 35.9 | 37.6 | 29.2 |
| 5. Was the science fair project required by your school? | Yes | 71.7 | 64.5 | 67.5 |
|  | No | 14.9 | 18.2 | 16.8 |
|  | Satisfied School Project | 12.0 | 14.3 | 15.2 |
| 6. Do you think science fair projects should be optional or required? (Need not be for competition.) | Optional | 74.1 | 75.4 | 74.4 |
|  | Required | 25.1 | 23.1 | 25.6 |
| 8. Do you think science fair projects for competition should be optional or required? | Optional | 83.8 | 83.9 | 78.8 |
|  | Required | 14.4 | 13.6 | 20.9 |
| 10. From whom do you think it would be reasonable to receive help? | 1. Parents | 77.0 | 77.4 | 78.0 |
|  | 2. Siblings | 63.1 | 61.2 | 61.4 |
|  | 3. Other family members | 57.6 | 54.3 | 52.9 |
|  | 4. Teachers | 90.3 | 90.7 | 89.8 |
|  | 5. Other students | 64.1 | 61.6 | 61.2 |
|  | 6. Scientists | 70.7 | 67.7 | 70.0 |
|  | 7. A paid mentor | 33.0 | 27.5 | 27.0 |
|  | 8. Articles on the Internet | 78.8 | 77.1 | 78.2 |
|  | Articles in books or magazines | 73.3 | 70.6 | 72.5 |
|  | Other | 3.9 | 2.3 | 3.9 |
| 11. Who actually helped you? | 1. Parents | 46.3 | 48.2 | 51.0 |
|  | 2. Siblings | 16.5 | 10.8 | 14.3 |
|  | 3. Other family members | 4.5 | 5.4 | 7.7 |
|  | 4. Teachers | 53.7 | 51.8 | 55.4 |
|  | 5. Other students | 30.1 | 30.7 | 23.7 |
|  | 6. Scientists | 8.6 | 8.0 | 8.0 |
|  | 7. A paid mentor | 0.5 | 0.4 | 1.1 |
|  | 8. Articles on the Internet | 52.6 | 57.5 | 57.9 |
|  | Articles in books or magazines | 23.3 | 22.7 | 24.0 |
|  | Other | 5.8 | 3.4 | 5.2 |
| 12. Kind of help reasonable to expect from others? | 1. Being given the main idea | 18.3 | 18.9 | 21.2 |
|  | 2. Development of the idea | 39.0 | 45.6 | 40.5 |
|  | 3. Gathering background research information, or finding a research site or participants | 46.1 | 45.0 | 49.3 |
|  | 4. Performing the experiments | 43.2 | 46.2 | 43.5 |
|  | 5. Writing the report | 12.6 | 14.7 | 13.5 |
|  | 6. Fine tuning the report after it is written | 58.9 | 62.4 | 52.3 |
|  | 7. Designing the poster board and presentation | 33.5 | 35.9 | 33.1 |
|  | 8. Producing charts or graphs | 23.0 | 24.4 | 19.3 |
|  | 9. Coaching for the interview with judges | 63.1 | 59.1 | 58.4 |
|  | 10. Copying the project from someone else | 1.6 | 2.7 | 1.4 |
|  | Other | 2.4 | 2.1 | 3.3 |
| 13. What kind of help did you actually receive? | 1. Being given the main idea | 8.9 | 9.4 | 13.8 |
|  | 2. Development of the idea | 25.7 | 26.8 | 26.2 |
|  | 3. Gathering background research information, or finding a research site or participants | 27.2 | 25.8 | 25.9 |
|  | 4. Performing the experiments | 29.3 | 28.6 | 24.5 |
|  | 5. Writing the report | 6.8 | 9.5 | 8.5 |
|  | 6. Fine tuning the report after it is written | 35.9 | 33.2 | 32.2 |
|  | 7. Designing the poster board and presentation | 19.9 | 22.3 | 19.0 |
|  | 8. Producing charts or graphs | 11.8 | 13.9 | 7.7 |
|  | 9. Coaching for the interview with judges | 19.4 | 20.9 | 22.0 |
|  | 10. Copying the project from someone else | 0.3 | 0.7 | 1.1 |
|  | Other | 6.5 | 6.1 | 8.8 |
| 14. Did you get the kind of help you wanted from teachers? | Yes | 69.9 | 74.3 | 72.2 |
|  | No | 28.0 | 24.6 | 26.7 |
| 16. Did you get the amount of help you wanted from teachers? | Yes | 65.4 | 73.1 | 74.1 |
|  | No | 31.9 | 25.2 | 24.5 |
| 17. Were the results of your project as expected? | Yes | 63.6 | 65.6 | 63.1 |
|  | No | 34.8 | 32.8 | 35.5 |
| 18. What obstacles did you face? | 1. Coming up with the main idea | 47.9 | 45.6 | 44.6 |
|  | 2. Getting motivated to do the project | 46.1 | 43.1 | 35.5 |
|  | 3. Becoming disappointed with the project | 29.1 | 25.5 | 21.8 |
|  | 4. Limited resources | 39.8 | 38.1 | 36.1 |
|  | 5. Limited knowledge | 29.1 | 30.9 | 30.6 |
|  | 6. Limited skills | 23.6 | 24.5 | 22.0 |
|  | 7. Limited cooperation | 15.4 | 12.7 | 11.6 |
|  | 8. Getting organized | 23.3 | 26.8 | 21.2 |
|  | 9. Time pressure | 59.2 | 62.6 | 57.3 |
|  | 10. Not enough money | 21.5 | 16.7 | 18.7 |
|  | 11. Results not as expected | 24.6 | 22.8 | 20.7 |
|  | Other | 4.2 | 3.7 | 4.4 |
| 19. How did you overcome the obstacles? | 1. Used someone else's main idea | 1.8 | 1.3 | 2.8 |
|  | 2. Picked a familiar / interesting topic | 34.0 | 30.8 | 29.8 |
|  | 3. Did more background research | 49.7 | 50.4 | 48.5 |
|  | 4. Stopped working on the project for a while | 17.5 | 15.8 | 16.0 |
|  | 5. Made a timeline to follow | 19.9 | 22.3 | 20.7 |
|  | 6. Perseverance and self-discipline | 41.6 | 48.1 | 44.4 |
|  | 7. Had someone else to keep me on track | 16.0 | 16.2 | 13.8 |
|  | 8. Had someone else do the math | 1.3 | 1.6 | 1.1 |
|  | 9. Changed the research plan | 14.7 | 18.7 | 11.6 |
|  | 10. Collected more data | 23.6 | 24.6 | 22.0 |
|  | 11. Had someone else collect the data | 1.0 | 2.1 | 0.6 |
|  | 12. Used someone else's data | 0.3 | 0.5 | 1.4 |
|  | 13. Made up the data | 5.0 | 4.1 | 4.1 |
|  | 14. Changed the hypothesis to fit the data | 4.7 | 3.8 | 4.4 |
|  | 15. Changed the data to fit the hypothesis | 3.1 | 1.5 | 2.2 |
|  | Other | 6.0 | 4.5 | 8.0 |
| 20. Are you interested in a career in the sciences or engineering? | Yes | 53.9 | 55.2 | 57.6 |
|  | No | 17.5 | 19.0 | 15.2 |
|  | Not Sure | 28.0 | 25.6 | 27.3 |
| 21. Did your science fair experience increase your interest in the sciences or engineering? | Yes | 52.4 | 55.4 | 58.7 |
|  | No | 46.9 | 44.1 | 40.8 |

**Supplemental Table 2.** Factors influencing the effect of SEF participation on Asian students’ interest in S&E

| Survey Questions | Answers | SEF increased my interest in S&E |  |  |
| --- | --- | --- | --- | --- |
|  |  | Yes % (#)<br>(276 students) | No % (#)<br>(162 students) | P value |
| Interested in a career in S&E | Yes | 77.2 (213) | 41.4 (67) | <.001 |
| Level of SEF competition? | District, Region or State | 49.3 (136) | 17.9 (29) | <.001 |
| SEF required? | Yes | 42.8 (118) | 80.2 (130) | <.001 |
| Project Team or Individual? | Individual | 65.6 (181) | 42.6 (69) | <.001 |
| Participation? | Did SEF > once | 33.0 (91) | 32.1 (52) | .851 |
| Who helped with your SEF project? (more than one answer is possible) | Parents | 48.6 (134) | 43.8 (71) | .338 |
|  | Teachers | 59.8 (165) | 43.2 (70) | <.001 |
|  | Scientists | 14.1 (39) | 3.7 (6) | <.001 |
|  | Articles on the internet | 63.4 (175) | 54.9 (89) | .080 |
|  | Articles in books or magazines | 32.2 (89) | 15.4 (25) | <.001 |
| Received kind of help needed from teachers? | Yes | 80.8 (223) | 55.6 (90) | <.001 |
| Types of help received? | Gathering background info, research site and participants | 29.0 (80) | 23.5 (38) | .208 |
|  | Fine tuning the report | 36.2 (100) | 23.5 (38) | .005 |
|  | Coaching for the interview | 33.0 (91) | 9.3 (15) | <.001 |
| Obstacles faced? | Getting organized | 23.2 (64) | 27.2 (44) | .352 |
|  | Time Pressure | 67.4 (186) | 65.4 (106) | .675 |
| Ways to overcome obstacles? | More background research | 61.6 (170) | 45.7 (74) | .001 |
|  | Made a timeline | 31.9 (88) | 16.0 (26) | <.001 |
|  | Perseverance | 55.4 (153) | 38.3 (62) | <.001 |

**Supplemental Table 3.**
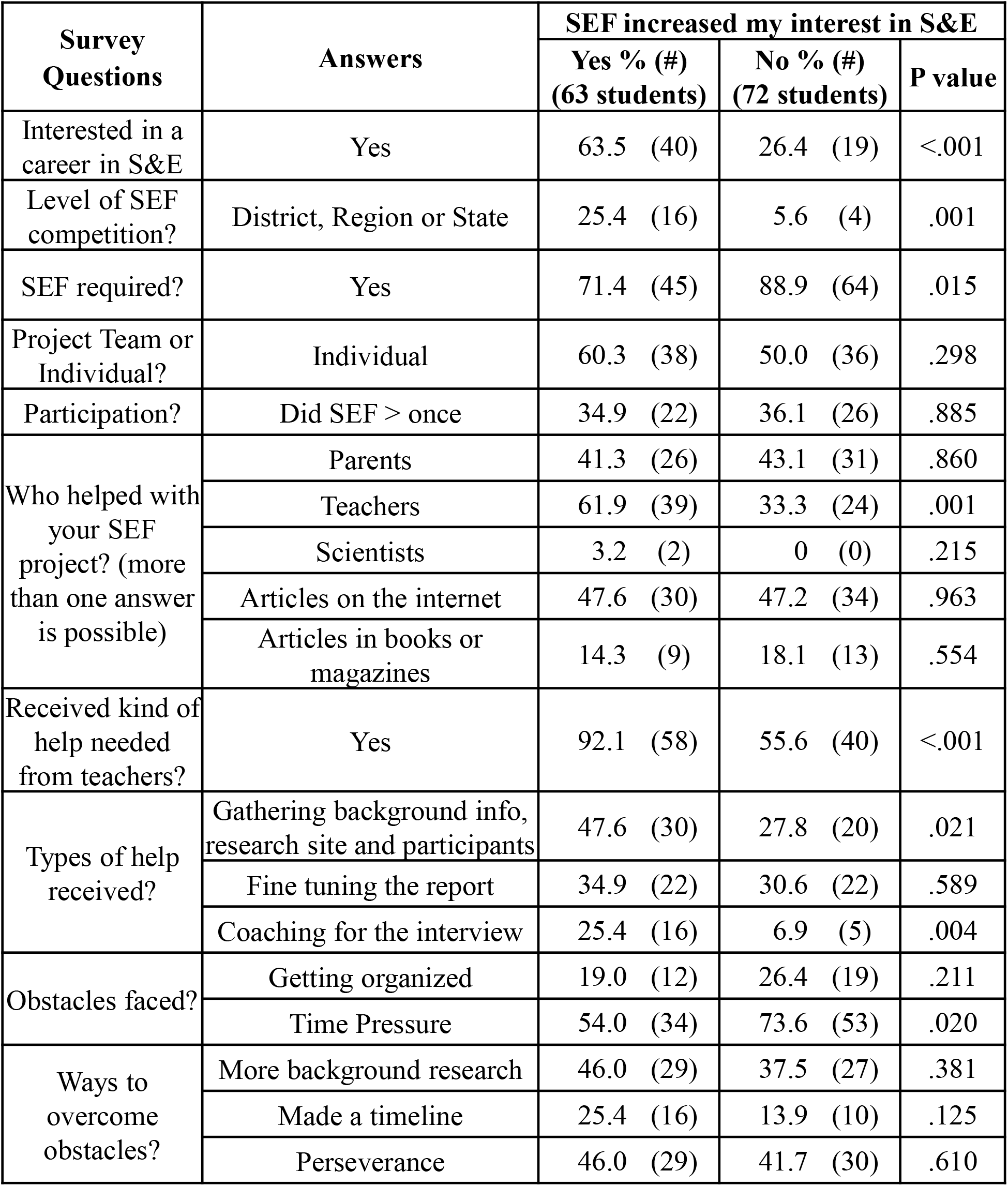
Factors influencing the effect of SEF participation on black students’ interest in S&E

**Supplemental Table 4.** Factors influencing the effect of SEF participation on Hispanic students’ interest in S&E

| Survey Questions | Answers | SEF increased my interest in S&E |  |  |
| --- | --- | --- | --- | --- |
|  |  | Yes % (#)<br>(159 students) | No % (#)<br>(90 students) | P value |
| Interested in a career in S&E | Yes | 67.9 (108) | 37.8 (34) | <.001 |
| Level of SEF competition? | District, Region or State | 64.8 (103) | 27.8 (25) | <.001 |
| SEF required? | Yes | 59.1 (94) | 77.8 (70) | .003 |
| Project Team or Individual? | Individual | 66.7 (106) | 56.7 (51) | .133 |
| Participation? | Did SEF > once | 49.1 (78) | 27.8 (25) | .001 |
| Who helped with your SEF project? (more than one answer is possible) | Parents | 47.8 (76) | 28.9 (26) | .005 |
|  | Teachers | 62.9 (100) | 38.9 (35) | <.001 |
|  | Scientists | 13.8 (22) | 4.4 (4) | .029 |
|  | Articles on the internet | 62.9 (100) | 47.8 (43) | .024 |
|  | Articles in books or magazines | 28.3 (45) | 16.7 (15) | .045 |
| Received kind of help needed from teachers? | Yes | 79.2 (126) | 67.8 (61) | .049 |
| Types of help received? | Gathering background info, research site and participants | 25.2 (40) | 34.4 (31) | .144 |
|  | Fine tuning the report | 33.3 (53) | 18.9 (17) | .019 |
|  | Coaching for the interview | 28.9 (46) | 8.9 (8) | <.001 |
| Obstacles faced? | Getting organized | 26.4 (42) | 32.2 (29) | .330 |
|  | Time Pressure | 62.3 (99) | 54.4 (49) | .227 |
| Ways to overcome obstacles? | More background research | 62.9 (100) | 36.7 (33) | <.001 |
|  | Made a timeline | 24.5 (39) | 5.6 (5) | <.001 |
|  | Perseverance | 5.9 (81) | 3.0 (27) | .001 |

**Supplemental Table 5.** Factors influencing the effect of SEF participation on white students’ interest in S&E

| Survey Questions | Answers | SEF increased my interest in S&E |  |  |
| --- | --- | --- | --- | --- |
|  |  | Yes % (#)<br>(198 students) | No % (#)<br>(236 students) | P value |
| Interested in a career in S&E | Yes | 73.2 (145) | 31.4 (74) | <.001 |
| Level of SEF competition? | District, Region or State | 40.4 (80) | 17.8 (42) | <.001 |
| SEF required? | Yes | 63.1 (125) | 80.5 (190) | <.001 |
| Project Team or Individual? | Individual | 70.2 (139) | 56.8 (134) | .005 |
| Participation? | Did SEF > once | 35.9 (71) | 34.3 (81) | .738 |
| Who helped with your SEF project? (more than one answer is possible) | Parents | 56.1 (111) | 52.1 (123) | .411 |
|  | Teachers | 60.6 (120) | 46.2 (109) | .003 |
|  | Scientists | 12.1 (24) | 3.0 (7) | <.001 |
|  | Articles on the internet | 62.1 (123) | 50.0 (118) | .012 |
|  | Articles in books or magazines | 25.3 (50) | 19.1 (45) | .121 |
| Received kind of help needed from teachers? | Yes | 84.3 (167) | 67.8 (160) | <.001 |
| Types of help received? | Gathering background info, research site and participants | 25.3 (50) | 19.1 (45) | .121 |
|  | Fine tuning the report | 47.5 (94) | 36.0 (85) | .019 |
|  | Coaching for the interview | 22.2 (44) | 14.8 (35) | .047 |
| Obstacles faced? | Getting organized | 17.2 (34) | 33.1 (78) | <.001 |
|  | Time Pressure | 58.1 (115) | 58.5 (138) | .934 |
| Ways to overcome obstacles? | More background research | 56.1 (111) | 39.8 (94) | .001 |
|  | Made a timeline | 26.3 (52) | 16.9 (40) | .019 |
|  | Perseverance | 50.0 (99) | 42.8 (101) | .134 |

